# A Compact Base Editor Rescues AATD-associated Liver and Lung Disease in Mouse Models

**DOI:** 10.1101/2025.05.07.652636

**Authors:** Jenny Gao, Nathan Bamidele, Debora Pires-Ferreira, April Destefano, Qiushi Tang, Yueying Cao, Jun Xie, Guangping Gao, Alisha Gruntman, Erik Sontheimer, Terence R. Flotte, Wen Xue

## Abstract

Alpha-1 antitrypsin deficiency (AATD) is commonly caused by a G-to-A mutation in the *SERPINA1* gene (the PiZ mutation). The mutant PiZ AAT protein is sequestered in hepatocytes, causing lung emphysema due to insufficient AAT protein to inhibit neutrophil elastase in the lung. Here we show that a compact adenine base editor (ABE) with an evolved Cas9 nickase derived from *Neisseria meningitidis* (eNme2.C) can be packaged in a single AAV and correct the PiZ mutation in mouse models of AATD. An all-in-one eNme2.C-TadA8e/guide 2 plasmid achieved approximately 20% on-target editing in PiZ reporter cells. TadA9e, which has a narrower editing window than TadA8e, reduced bystander editing without significantly affecting the on-target edit. In PiZ transgenic mice, eNme2.C-TadA9e AAV showed approximately 23% editing efficiency after 8 weeks and reduced liver disease burden in treated mice. In a new AAT-null;PiZ transgenic mouse model, ABE restored serum levels of AAT to beyond the 570 μg/mL therapeutic level. Moreover, ABE treatment was able to significantly correct lung functions in AAT-null;PiZ animals with emphysema. This study demonstrates the feasibility of an eNme2.C-based ABE in a single AAV to treat both AATD-associated liver and lung disease.

## Introduction

Alpha-1 antitrypsin deficiency (AATD) is an inherited monogenic disorder that can lead to lung disease and liver damage^1–3^. It is caused by mutations in the SERine Proteinase Inhibitor family A member 1 (*SERPINA1)* gene, which encodes AAT—a serine protease inhibitor made by hepatocytes and secreted into serum for delivery to the lungs, where it predominantly inhibits neutrophil elastase (NE). The most common variant is known as the Z allele (PiZ). The PiZ allele is a G-to-A mutation that causes a glutamic acid to lysine amino acid change at position 342 of AAT (Z-AAT). The Z-AAT protein misfolds, polymerizes, and accumulates in the liver, increasing the risk of liver fibrosis^4–6^. The reduction in serum AAT levels leads to lung emphysema due to the destruction of alveolar spaces by uninhibited neutrophil elastase. Currently, the only approved therapy for AATD-associated emphysema is costly, lifelong AAT infusions. AATD exhibits a remarkably strong founder effect in Europe and North America. In fact, greater than 95% of patients with severe AATD lung disease possess at least one PiZ allele^7^. This unusual genetic homogeneity of the AATD population makes it uniquely suitable for a base editing strategy.

Traditional gene therapies are being developed to augment AAT function in patients. Strategies focus on the delivery of adeno-associated virus (AAV) vectors encoding a wild-type *SERPINA1* transgene or modulating RNA levels of the mutant transcript^8–18^. However, these approaches still require repeated treatment or do not reach the therapeutic threshold needed to ameliorate disease phenotype. A one-time treatment that corrects the *SERPINA1* gene and halts AATD progression can significantly improve patient quality of life.

Base editors (BE) have revolutionized gene therapy treatments due to their ability to make precise edits^19–33^. Currently, the two major classes of BEs are cytosine base editors (CBEs) which make C-to-T edits^28^, and adenine base editors which make A-to-G edits (ABEs)^22^. Despite their promise, BEs have been challenging to deliver with a single AAV due to the packaging limit (∼4.7kb cargo size) that must encapsulate the BE, guide cassette, promoters, and regulatory elements^34, 35^. The most commonly used Cas9 is from *Streptococcus pyogenes* (SpCas9), which alone is 1368 amino acids, and hence is unable to be packaged into a single AAV as an ABE system. AAVs have been widely used as a gene therapy tool in clinics, as they have low integration risk, low immunogenicity, and can target a myriad of tissues by switching the virus serotype^8, 36–43^. However, its packaging limit stands at ∼4.7kb. Given that base editors with the most widely used *Streptococcus pyogenes* Cas9 (SpCas9) are already ∼5.2kb without the guide and regulatory sequences, we sought to use other editors. A single AAV can minimize AAV dosage, thereby reducing potential toxicity and increasing editing efficiency since cells only need a single vector for editing to occur.

In this study, we used an evolved *Neisseria meningitidis* (eNme2) Cas9-based ABE system with a single C PAM to correct the pathogenic PiZ mutation using a single AAV^36, 44–49^. We then optimized for low bystander edits and high on-target edits by changing the TadA deaminase domain. The all-in-one ABE was then packaged into AAV and injected into animals carrying the human PiZ allele. Treated animals showed correction of the PiZ mutation, along with reduced liver disease burden. Moreover, total serum AAT levels increased beyond 570 μg/mL and emphysema improved in treated animals.

## Results

### Optimizing ABE for low bystander edits and high on-target edit

To edit the PiZ site, we aimed to develop a compact ABE system that can be packaged into a single adeno-associated virus (AAV). We started with the evolved *Neisseria meningitidis* Cas9 (eNme2.C)^47^, which has been shown to have high base editing capabilities. Moreover, it is ∼3.2kb, thus small enough to be packaged into a single AAV for therapeutic delivery. First, we tested two guides compatible with an eNme2.C Cas9 ABE system, which uses the N_4_C PAM. We designed two sgRNAs (guide 1 or 2) with slightly offset editing windows^47^ **(Fig. 1a, 1b**). To study PiZ allele editing events by ABEs, we transduced HEK293T cells with a lentiviral construct with the coding sequence of the Z allele, generating a PiZ reporter line. We transfected PiZ reporter cells with constructs encoding eNme2.C-TadA8e and one of two sgRNAs (guide 1 or 2) and performed high-throughput sequencing using primers specific to the reporter that cannot amplify the endogenous *SERPINA1* gene (**Supplemental file 4**). For guide 2, the target A in PiZ is at position 9 (A9). To directly compare the editing rate, we defined the same “A” positions for guides 1 and 2. Guides 1 and 2 showed 36.98% and 33.84% editing, respectively (**Fig. 1c**). Although guide 1 showed slightly higher on-target editing at the A9 site, the bystander rates were equally high – bystander editing ranged from 0.03% to 33.26% compared to the 0.03% to 12.19% with guide 2. Given the lower bystander rate but comparable on-target editing rate of guide 2, we chose guide 2 for all subsequent experiments.

**Figure 1.**
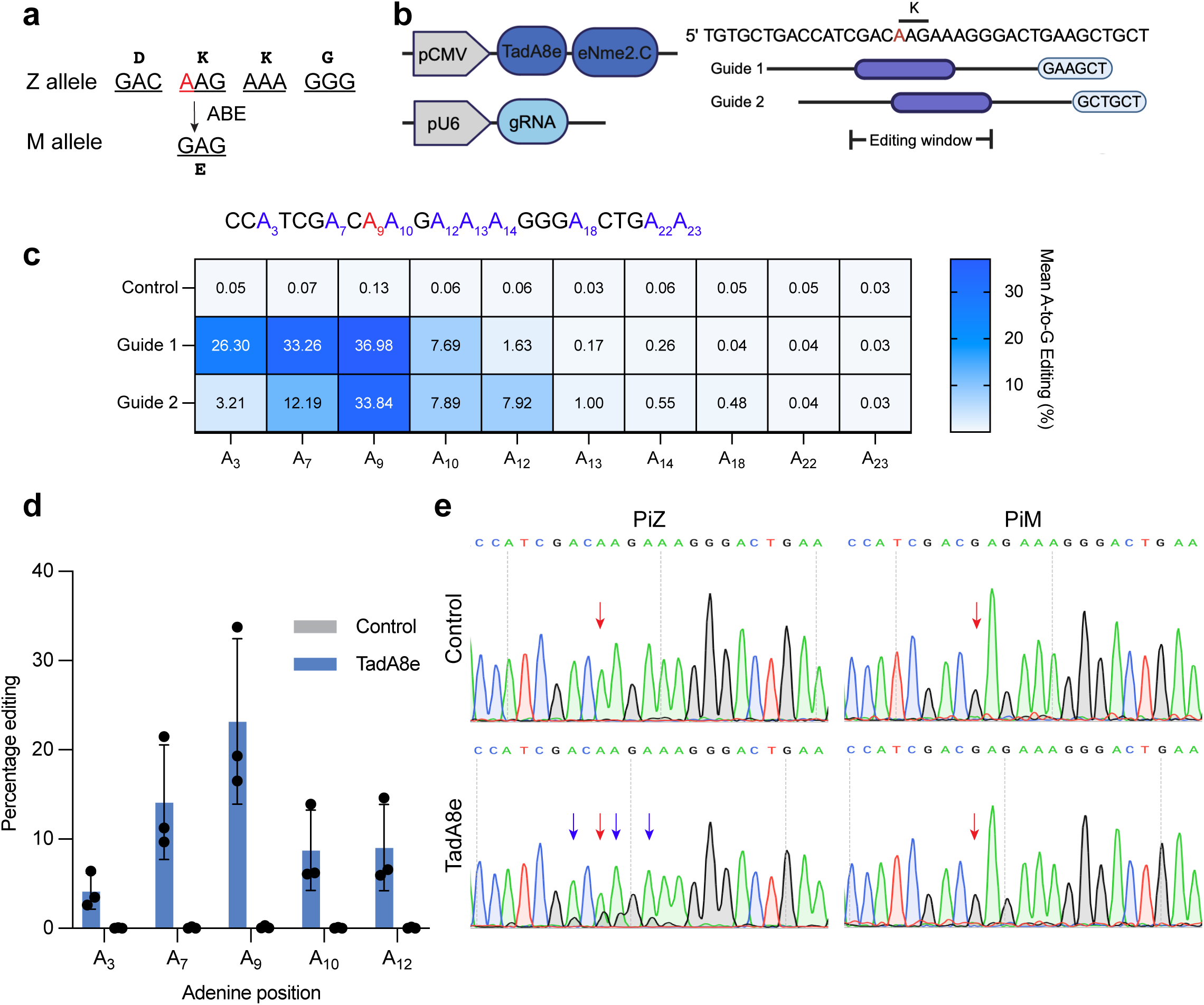
Optimization of an all-in-one editor for AAT correction. (a) PiZ mutation site with the pathogenic A mutation noted in red. Amino acids are in bold. (b) Two guides were tested and delivered to PiZ reporter cells. PAM and editing window for each guide are shown. (c) A to G editing efficiency of the two guides in PiZ reporter cells. Numbers represent mean, n=3. Target A is in red, with potential bystander edits in blue. (d) Editing efficiency of all-in-one construct that can deliver eNme2.C-TadA8e. Mean ± SD shown, n=3. (e) Mutant-specific sgRNA edits PiZ reporter but not the wildtype PiM with a single G mismatch. Red arrow depicts target base while blue depicts bystanders. Representative image shown.

We cloned eNme2.C and guide 2 into a single vector (hereby called eNme2.C-TadA8e) driven by the U1a promoter. eNme2.C-TadA8e showed an average of 23.18% on-target editing at the A9 position, with bystander rates ranging from 4.16% to 14.13% (**Fig. 1d**).

Although some bystander edits have been characterized as not affecting AAT protein function or serum secretion, we aimed to reduce these bystander edits further, as they can generate mutant AAT variants with unknown function and cause an unknown immune response^13^. TadA9e is a derivative of TadA8e but with a narrower editing window due to weaker TadA-DNA interaction^32, 50^. To test its activity, we swapped the TadA8e domain with TadA9e, thereby generating the all-in-one eNme2.C-TadA9e. This all-in-one vector proved to have efficient in vitro editing, with on-target editing averaging 17.94% (**Fig. 2a**). When bystander editing was normalized to on-target A9 editing, eNme2.C-TadA9e showed lower bystander editing compared to eNme2.C-TadA8e and had more perfect alleles (**Fig. 2b, 2c**). Thus, we used the all-in-one eNme2.C-TadA9e for all subsequent experiments.

**Figure 2.**
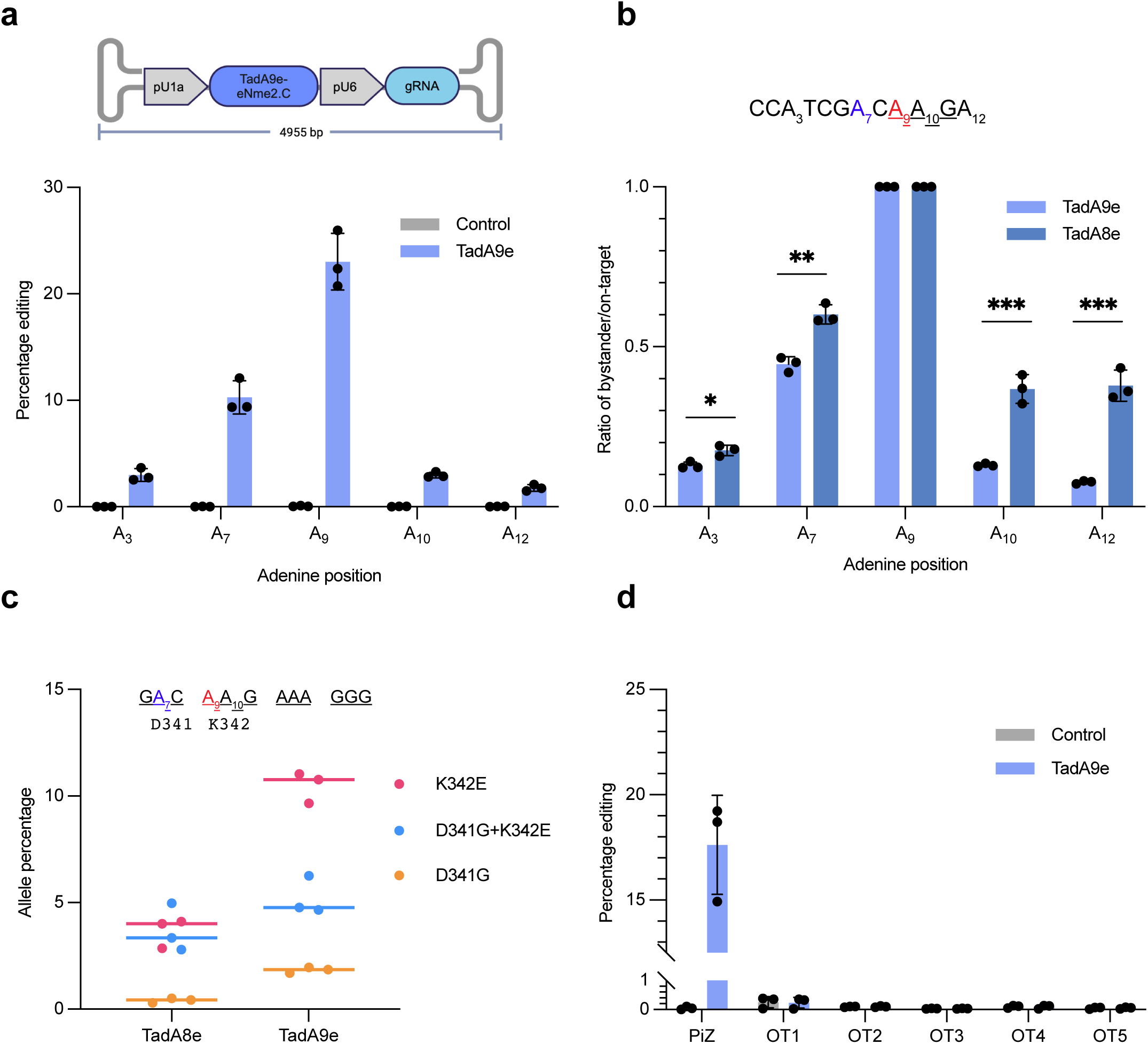
Reduction of bystander edits with TadA9e. (a) Editing efficiency of all-in-one construct that can deliver eNme2.C-TadA9e in reporter cells. Mean ± SD shown, n=3. (b) Ratio of bystander to on-target edit in PiZ reporter cells. Mean ± SD shown. *p ≤ 0.05, **p ≤ 0.01,***p ≤ 0.001 by unpaired t-test. (c) Allele population of cells edited with eNme2.C-TadA8e or eNme2.C-TadA9e. Line depicts median, n=3. (d) Deep sequencing of OT sites. Mean ± SD shown. See Supplemental file 1 for additional information, n=3.

### Assessment of eNme2.C ABE off-target edits

Nme2-based editors have higher fidelity compared to the more widely used SpyCas9^44, 47, 49^. Despite this, we aimed to characterize the off-target editing of eNme2.C-TadA9e through computational and experimental approaches. First, we sequenced the endogenous wild-type PiM site in the PiZ reporter cell transfected with the all-in-one eNme2.C-TadA8e to detect any editing. The absence of any editing at the PiM site, which differs from the PiZ site by a single nucleotide, points to the high fidelity of our system (**Fig. 1e**).

Cas9-based editing modalities can induce both Cas-independent and -dependent off-target edits^51^. Cas-dependent off-target editing is due to sequence similarities between the target protospacer and off-target protospacer, allowing the guide to bind and hence editing to occur^52–57^. Cas-independent off-targeting editing is due to long-term expression of the TadA deaminase enzyme, causing low-level deamination across the genome and transcriptome^58–63^. We sought to characterize the Cas-dependent off-targets with Cas-OFFinder^64^. The Cas-OFFinder algorithm nominated 150 sites with five or fewer mismatches to the target protospacer. After filtering for suitable PAM sites, and sites with conserved seed regions, defined as nucleotides 17-23^49^, we tested the editing rate at five sites by high-throughput sequencing (**Supplemental file 1**). There was no difference in editing between the control and transfected cells (**Fig. 2d**).

### All-in-one eNme2.C-TadA9e treated animals have significant in vivo editing with decreased liver disease burden

We then sought to test the all-in-one eNme2.C-TadA9e in PiZ animals, which carries the human PiZ transgene integrated^4^. The vector was packaged into AAV8, and 9e11 vector genomes (vg) was delivered to animals via tail-vein injection. Control animals were left untreated. After 4 or 8 weeks, the liver of each animal was collected, and the PiZ site was amplified for targeted amplicon sequencing (**Fig. 3a**). At 4 weeks, on-target editing was 16.75% and this increased to 22.83% at 8 weeks (**Fig. 3b**). Allele analysis showed that in the treated animals, perfect editing (K342E allele, A9G) represented 12.62% of the allele population at 4 weeks, and 16.53% at 8 weeks (**Fig. 3c**). The D341G + K342E (A7G+A9G) allele and D341G (A7G) allele were the other top alleles, representing 1.62% and 2.08%, respectively, of the allele population at 8 weeks (**Fig. 3c**). These data show that the perfectly edited allele presents the majority of the edited alleles.

**Figure 3.**
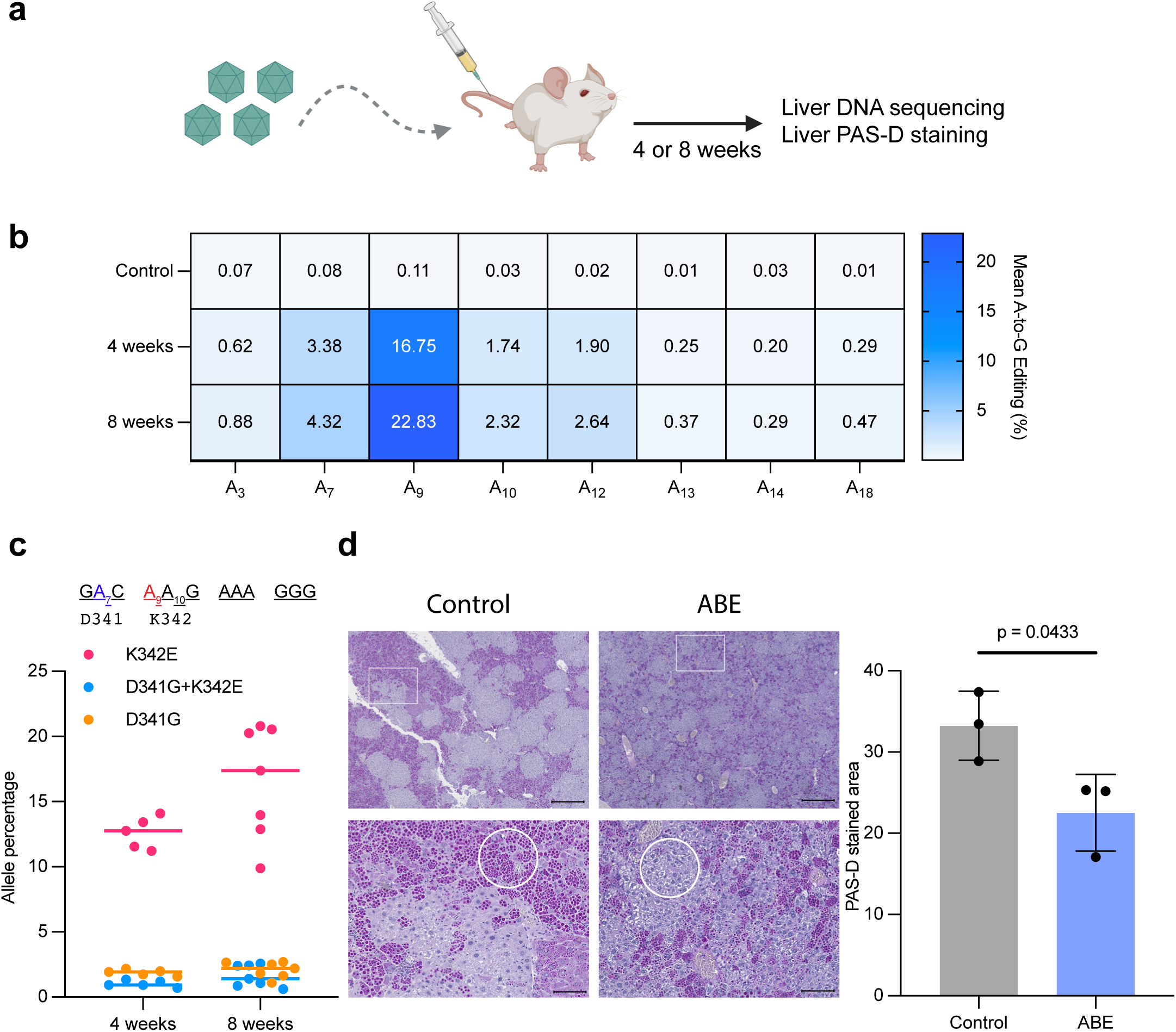
In vivo editing of PiZ animals. (a) Schematic of tail-vein injection in PiZ animals with all-in-one eNme2.C-TadA9e. (b) eNme2.C-TadA9e in vivo editing at 4 and 8 weeks in PiZ animals. Numbers represent mean (n=3 for control, n=5 for 4 weeks, and n=7 for 8 weeks). (c) Allele percentage in PiZ mice treated with eNme2.C-TadA9e. Line represents median, n=5 for 4 weeks, n=7 for 8 weeks. (d) Representative images of mouse liver stained with PAS-D. Mice were either untreated or treated with 9e11vg AAV for 8 weeks. Scalebar: 500µm for top images, 100µm for bottom images. Squares represent area of zoom from the top images, while circle represents the PASD+ and PAS-regions. Right: quantification. Mean ± SD shown. Significance was calculated by unpaired t-test, n=3.

It was previously reported that significant base editing correction of the PiZ site can lead to decreased AAT globules in the liver^13, 17^, which can be detected with Periodic acid–Schiff–diastase stain (PAS-D stain). We sought to investigate if our editing rate was sufficient and performed PAS-D staining on mouse liver. Indeed, animals treated for 8 weeks showed a significant decrease in PAS-D positive area (p=0.04) (**Fig. 3d**).

### Generation of a clinically relevant Aat-null;PiZ transgenic mouse model

The PiZ animal carries multiple copies of the human Z-AAT allele, along with five isoforms of the murine *Serpina1* gene. Due to the presence of its endogenous *Serpina1* genes, the PiZ animal does not develop emphysema. The five copies of the *Serpina1* gene were knocked out in C57BL/6J animals to generate AAT-null mice, which develop emphysema both spontaneously and through LPS challenges^65, 66^. To study the therapeutic advantage of ABE for correcting human Z-AAT, we crossed the AAT-null animals with PiZ animals to generate AAT-null;PiZ (**Fig. 4a**). Given that this strain carries multiple copies of the human Z-AAT allele, but no copies of murine *Serpina1*, it will develop both liver and lung disease.

**Figure 4.**
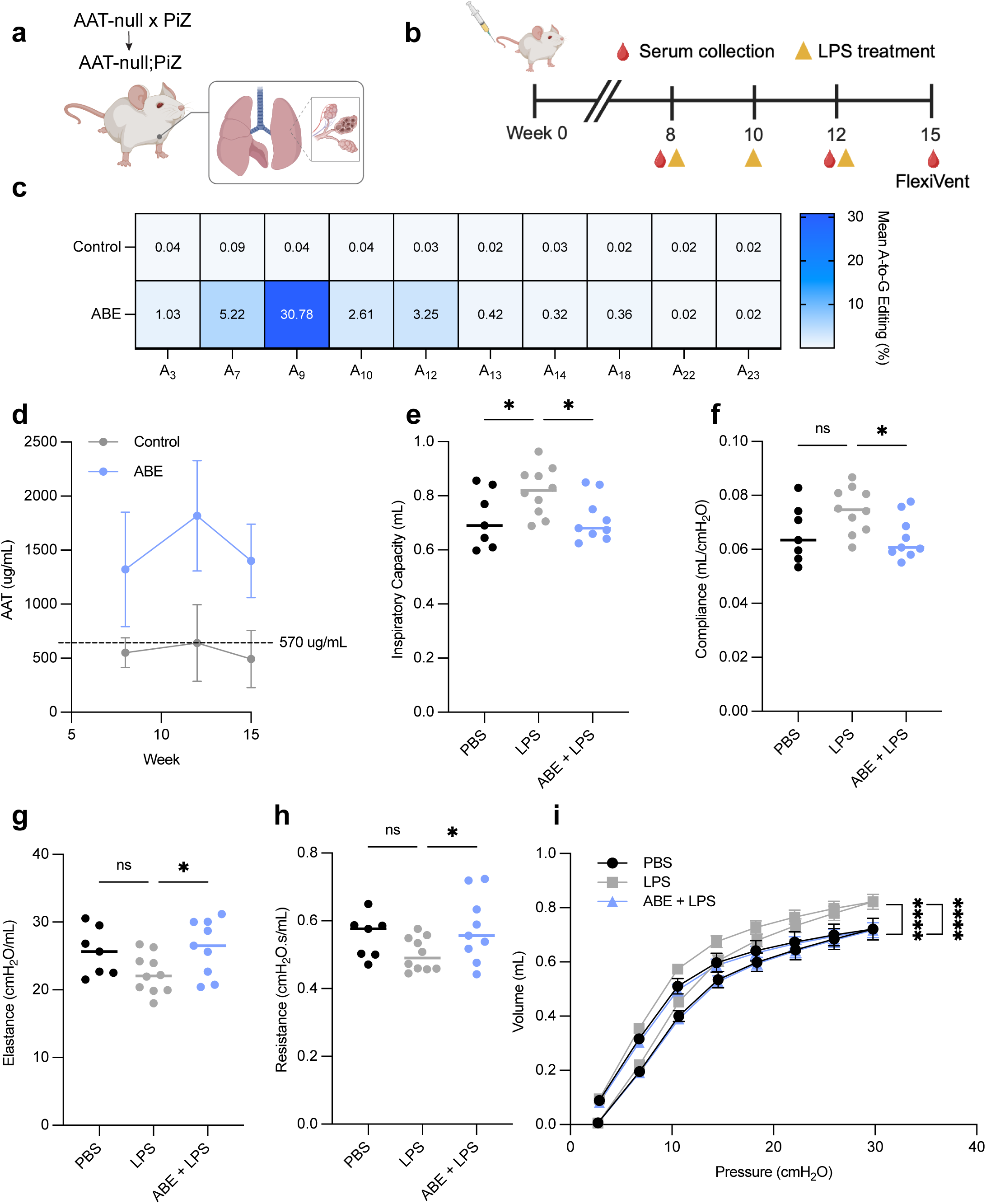
Lung disease reversal in AAT-null;PiZ animal. (a) Schematic of AAT-null;PiZ mouse generation (b) Schematic of ABE treatment to study LPS-induced emphysema in AAT-null;PiZ animals. (c) Editing in AAT-null;PIZ animals when treated with 9e11vg AAV for 15 weeks (n=4 for control, n=6 for ABE) (d) AAT levels in control and treated AAT-null;PiZ animals. Mean ± SD shown, n=3 for control, n=5 for ABE. Both groups were treated with LPS. (e-i) Inspiratory capacity (e), compliance (f), elastance (g), resistance (h) and (i) pressure-volume (PV) loop of PBS, LPS, or ABE+LPS treated animals (n=7, n=10, and n=9, respectively). Line depicts median. Significance was detected by one-way ANOVA with uncorrected Fisher’s least significance difference, except for PV-loop (two-way ANOVA with Dunnett. Mean ± SEM shown), *p ≤ 0.05,****p ≤ 0.0001.

We aimed to test the editing efficiency of our system in the AAT-null;PiZ animal. Animals were injected with 9e11vg AAV and followed for 15 weeks. Treated animals showed 30.78% on-target editing at 15 weeks (**Fig. 4c**) with perfect allele reaching 23.10% (**Supplemental Fig. S1**). Serum was collected from animals at 8, 12, and 15 weeks to test for human AAT. It is estimated that serum AAT level below 11 μM (570 μg/mL) significantly increases the risk of emphysema development. Because there was no suitable M-AAT specific antibody for ELISA, we used an antibody recognizing both Z- and M-AAT. In treated animals, total AAT levels increased to an average of 1,321.70 μg/mL at 8 weeks and consistently stayed above 570 μg/mL throughout the 15-week timeline (**Fig. 4d, Supplemental file 2**). To estimate the level of edited AAT, the majority of which likely is the M-AAT wild-type variant, we subtracted the AAT levels from the untreated animals, who only express the Z-AAT protein. At 15 weeks, the differential AAT level in treated animals subtracted from control was 909 μg/mL on average (**Supplemental Fig. S2**).

### Reversal of the lung disease phenotype

AAT-null mice have been shown to manifest lung disease with pulmonary physiology consistent with an emphysema phenotype, but only after prolonged aging or exposure to a proinflammatory stimulus or tobacco smoke^65, 66^. Proinflammatory stimuli are thought to accelerate the development of emphysema in the AAT deficient state by recruiting neutrophils to the lung, triggering their release of NE in proximity to pulmonary interstitial elastin fibers. Without sufficient AAT to counteract NE, destruction of the extracellular matrix of the lung leads to hyperinflation (increased inspiratory capacity) and decreased pulmonary elastic recoil, with a resultant decrease in elastance and lung tissue resistance, and an increase in lung compliance - a picture that matches experimentally induced emphysema in mice^65, 67^. To demonstrate the therapeutic effect of base editing on the lung disease phenotype, pulmonary physiology was assessed with the flexiVent system in AAT-null;PiZ animals with or without LPS challenge, and in the base-edited mice with LPS challenge. The LPS challenge induced the expected emphysema physiology, consisting of increased inspiratory capacity (IC), increased static compliance (Cst), decreased elastance of the respiratory system (Ers), decreased respiratory system resistance (Rrs) and upwards shift of PV-loop (**Fig. 4e-i**). Animals treated with ABE for 8 weeks (at which serum AAT reached 1,321.70 μg/mL) prior to LPS treatment had significant improvement in lung emphysema. Compared to the LPS group, the ABE+LPS group had reduced IC, reduced Cst, increased Ers, and increased Rrs (**Fig. 4e-h**). Moreover, the PV loop shifted downwards, suggesting that the ABE treatment was able to significantly mitigate the effects of LPS-induced emphysema (**Fig. 4i**).

## Discussion

Alpha-1 antitrypsin deficiency affects approximately 100,000 people in the United States every year^68^. Although AAT-specific therapies, such as AAT protein infusion have been clinically successful, they still have specific drawbacks such as the need for weekly, costly treatments. Moreover, the only FDA-approved treatment for AATD-associated liver disease is organ transplant. A one-time therapy targeting the genetic cause can address these drawbacks.

In our study, we evaluated the therapeutic potential of a novel editing agent, eNme2.C-TadA9e, for treating AATD caused by the PiZ allele. We optimized our ABE for minimal bystander and sufficient on-target editing and assessed the off-target editing rate of our system through both computational and experimental validation.

The lack of editing at the endogenous PiM site in our PiZ reporter cell line suggests that ABE would not cause bystander editing in patients with the Pi*MZ genotype. Our system proved to be highly efficient in editing the PiZ allele across two animal models carrying the human allele, with editing reaching 30.78% by week 15. Notably, the AAV dosage, 9e11vg or 4.5e13vg/kg, is significantly lower than typical doses in clinical trials^69^. Given that high AAV doses can cause toxicity^70–72^, the low dosage further underscores eNme2.C-TadA9e as a promising candidate for AATD therapy. We anticipate that this editing percentage will continue to improve, as cells with the PiM allele have a survival advantage over cells with the PiZ allele^73^, and the ABE system may continue to edit the PiZ site. Indeed, our observations support this, as editing has been steadily increasing from 16.75% to 30.78% by week 15. Importantly, treated animals have a significant reduction in AAT globule accumulation – a key factor in the development of AATD-associated liver fibrosis.

To study the effect of editing on AATD-associated lung emphysema, we developed a new AATD mouse model by crossing the PiZ and AAT-null strain. The AAT-null;PiZ strain was able to recapitulate emphysema through sequential LPS doses. We noted that AAT-null;PiZ animals treated with ABE for 8 weeks had a serum AAT level of ∼1,322 μg/mL, surpassing the 570 μg/mL threshold needed for decreased risk of lung emphysema development. Finally, we assessed the therapeutic potential of ABE for resolving AATD-associated emphysema. After establishing a model of emphysema with the AAT-null;PiZ animals, we treated the animals with ABE. ABE+LPS treated AAT-null;PiZ animals demonstrated a significant reduction in inspiratory capacity, compliance, and an increase in elastance and resistance. Cumulatively, this efficient low-dose AAV application, combined with a high editing rate and significant disease improvement, positions eNme2.C-TadA9e as a promising candidate for future studies.

Although we provided significant evidence for the use of eNme2.C-TadA9e as a therapy for AAT, future work could address some limitations. For one, due to the absence of a good M-AAT specific antibody, we were only able to characterize the total AAT in the mouse serum. Moreover, the PiZ and AAT-null;PiZ mouse is not a perfect animal model to study AAT. Both strains carry multiple copies of the PiZ allele, and the Z-AAT level can fluctuate depending on the number of PiZ copies. Finally, future study is needed to address whether the Nme2 or ABE domain induces immune response *in vivo*. Despite these limitations, this study demonstrates the feasibility of a compact eNme2.C ABE in a single AAV to correct the PiZ mutation *in vivo*.

## Methods

### Molecular cloning

Plasmids expressing sgAAT were constructed using Gibson assembly using a gene block expressing the guide cassette and Addgene plasmid 122091. Plasmid expressing the guide RNA was cloned by Gibson assembly of gblock with the guide cassette and BfuAI digested Addgene 122091. Plasmid expressing the all-in-one eNme2.C-TadA8e was constructed by Gibson assembly using Addgene plasmid 121507 as a backbone, PCR-amplification of Addgene plasmid 185667 to retrieve eNme2.C-TadA8e, gene block containing U1a promoter, and PCR-amplification of sgAAT plasmid. Plasmid expressing the all-in-one eNme2.C-TadA9e was cloned by Gibson assembly of gblock with TadA9e with backbone from the all-in-one eNme2.C-TadA8e plasmid.

### Cell culture

Cells were maintained in Dulbecco’s Modified Eagle’s Medium (DMEM) supplemented with 10% (v/v) fetal bovine serum (Gibco) and 1% (v/v) Penicillin/Streptomycin (Gibco). Cells were cultured at 37 °C with 5% CO_2_. HEK293T and 293fs cells were obtained from ATCC.

### Generation of PiZ reporter line

The coding sequence of hAAT was packaged into lentivirus. Briefly, 293fs cells were plated into 6 wells to reach 70-80% confluency the next day. On the day of transfection, delta 8.2, VSV-G, and H633 were combined and transfected into cells using LT1.

For transduction, HEK293T cells were seeded into 6-well plates. The next day, virus and polybrene were added. Puromycin was used to select transduced cells for 3 days. PiZ integration was verified through PCR amplification with primers and subsequent Sanger sequencing.

### Transient transfection

For transfection, cells were plated on 12-well plates at 100,000 cells per well. The next day, cells were transfected with Lipofectamine 3000 (Invitrogen). For screening guides and Nme2 variants, 250ng of the effector and 83ng of the sgRNA were transfected. For testing all-in-one constructs, 1000ng of the plasmid was transfected into the PiZ reporter cell line. 3 days after transfection, genomic DNA was collected with quick extraction buffer (Epicenter). The lysate was incubated at 65 °C for 15 minutes, and 98 °C for 5 minutes.

### Off-target analysis

Off-target analysis was performed with Cas-OFFinder using 5 or fewer mismatches relative to the intended target site. To determine potential off-target sites, we first filtered for sites with appropriate PAMs, followed by sites with conserved seed regions. Sites of known genes were then sequenced with amplicon sequencing. The highest editing rate at each site is reported.

### Amplicon sequencing and data analysis

Sequencing library preparation was done as previously described. For the first round of PCR, primers with Illumina forward and reverse adapters were used to amplify the gene of interest using either 1ul cell culture lysate or 100ng mouse DNA. PCR was performed with Phusion Flash High-Fidelity PCR Master Mix at 98 °C for 10 s, then 20 cycles of 98 °C for 1 s, 55 °C for 5 s, and 72 °C for 7s, followed by a final 72 °C extension for 1 min. 1ul of PCR product was then used for the second round of PCR in which a unique Illumina barcode was added to each sample. As before, PCR was performed at 98 °C for 1 s, 55 °C for 5 s, and 72 °C for 7s, followed by a final 72 °C extension for 1 min. PCR 2 products were purified by gel extraction using the QIAquick Gel Extraction Kit (Qiagen) and quantified by Qubit dsDNA HS Assay Kit (Thermo Fisher Scientific). The library was sequenced on an Illumina MiniSeq instrument following the manufacturer’s protocols. Sequencing reads were demultiplexed using bcl2fastq (Illumina). To quantify the frequency of precise editing and indels, CRISPResso2^74^ was run in base editor mode with ‘min_average_read_quality’ > 30. See supplemental file 3 for all sequencing primers.

The ratio of on-target to bystander edit was computed by: editing percentage at bystander adenine/editing percentage at target adenine.

### AAV production

AAV was produced, purified, and titered by the University of Massachusetts Gene Therapy Vector Core.

### Mouse experiments

All experimental procedures were approved by the Institutional Animal Care and Use Committee (IACUC) at the University of Massachusetts Medical School. Animals were housed in cages with access to food and water *ad libitum*, in rooms with controlled temperature and illumination (12 h/12 h light-dark cycle). AAV was administered via tail-vein injection for treated animals.

AAT-null/PiZ animals were generated by crossing the AAT-null and PiZ animals. The human AAT gene was verified via genomic DNA PCR (5’ TTG AGG AGC GAG AGG CAG TT, 5’ GAG GCG CTT GTC AGG AAG AT).

For the PiZ animal study, both male and female 4–8-week-old animals were used. For the AAT-null;PiZ animal study, both male and female animals were used for the PBS, LPS and ABE + LPS group (9-16 week-old, 13-18 week-old, and 5–10-week-old animals, respectively). No animals were excluded from the study. PiZ animals were taken down at either 4, or 8 weeks post AAV treatment while AAT-null;PiZ animals were taken down 15 weeks post AAV treatment. 5 liver biopsies were harvested per animal and genomic DNA was collected using DNeasy Blood & Tissue Kit (Qiagen, 69506). Editing efficiency was computed by taking the average across all five lobes. For histology, liver lobes were fixed in 10% neutral buffered formalin overnight before being embedded in paraffin. Fixed sections were stained with PAS-D to visualize AAT globules. PAS-D staining was analyzed using QuPath color thresholding.

### ELISA

Animals were bled via facial vein to retrieve serum and ELISA was performed as previously described^75^. To detect hAAT, clear flat-bottom immuno nonsterile 96-Well Plates (Thermo Fisher Scientific, 3855) were coated with goat anti-human AAT antibody (Bethyl Laboratories, A80-122A) overnight. The next morning, plates were blocked with Intercept PBS blocking buffer (Licorbio, 927-70003) followed by the addition of serially diluted samples and standard curves (Athens research, 16-16-011609) in triplicates. The plate was incubated for an hour, and goat anti-human AAT antibody (Bethyl Laboratories, A80-122P) was added for one hour. Between each step, the plate was washed with PBST. 3,3’,5,5’-tetramethylbenzidine (TMB) peroxidase (Thermo Fisher Scientific, 50-674-21) was added to the wells and the reaction was stopped by adding 0.18M sulfuric acid. The plate was read at wavelength 450nm.

### LPS treatments

Animals were either treated with AAV or untreated at week 0, followed by lipopolysaccharides (LPS) or PBS challenges at weeks 8, 10, and 12. For LPS challenges, mice were anesthetized with an i.p. dose of ketamine/xylazine mixture (90 mg/kg and 4.5 mg/kg, respectively), then intratracheally cannulated with a 22-gauge angiocatheter using a mouse intubating board. LPS isolated from *Escherichia coli* O55:B5 (Cayman Chemical, item number 19660, batch 0710200-1) were administered (30 μL, 1 μg/mL in PBS) via direct injection through the angiocatheter during inspiration. Following instillation, the animals received 2-3 ventilations of air to ensure the solution was properly inhaled. After the LPS challenge, the mice were allowed to recover and return to the colony.

### FlexiVent

First, animals were anesthetized with a ketamine/xylazine mixture (100 mg/kg and 10 mg/kg, i.p.). A tracheotomy was performed to insert an 18-gauge metal cannula, and the animal was connected to the machine. Pulmonary function was assessed through the Forced oscillation technique (FOT) performed using the flexiVent FX system (SCIREQ Inc., Montreal Qc, Canada). Mice were ventilated with a tidal volume of 10 mL/kg, a frequency of 150 breaths/min, an inspiratory to expiratory ratio of 2:3, and a positive end-expiratory pressure of 3 cmH_2_O. Then, two deep inflations were given to maximally inflate the lungs and standardize lung volume. Inspiratory capacity (IC), compliance (C), elastance (Ers), and resistance (Rrs) were calculated from volume, pressure and flow signals collected during the test. The flow signal is derived from the volume signal. Pressure-volume curve was generated using a ramp-style pressure-driven maneuver (PVr-P). Static compliance (Cst) was measured using the PV loop between the pressures of 3 –7 cmH_2_O. Each measurement was taken three times, and the average was used for further analysis.

### Statistical analysis

All experiments were done with 3 biological replicates. Statistical analyses were conducted using Graph Pad Prism 8 (Graph Pad Software).

### Data availability

The raw sequencing data will be deposited to the NCBI Sequence Read Archive. All raw data will be available from the corresponding authors upon request.

## Acknowledgments

We thank Y. Liu in the UMass Morphology Core for support and G. Cottle for mouse injections. We thank Lauren Shumate and Tingting Jiang for cloning the PiZ reporter lentivirus plasmid. This work was supported by grants from the National Institutes of Health (P01HL158506, R01CA275945, R01GM150279, and UH3HL147367), and Friedreich’s Ataxia Research Alliance. J.G. was supported by NIH F30HL176024.

## Author Contributions

J.G., N.B., and D.P.-F designed the experiments. J.G. performed and analyzed all the cell culture experiments. Q.T., Y.C., and A.D. generated and bred the animals. J.G., D.P.-F, and A.D. performed in vivo experiments using AAV vectors generated by J.X. J.G. analyzed all in vivo data. G.G., A.G, E.J.S, T.R.F., and W.X. oversaw the experiments. J.G. and W.X. wrote the manuscript with input from all co-authors.

## Declaration of interests

E.J.S. is a co-founder and Scientific Advisory Board member of Intellia Therapeutics and a Scientific Advisory Board member at Tessera Therapeutics. The University of Massachusetts Chan Medical School has filed patent applications related to this work. All other authors have no competing interests.

---

**Supplemetary Figure S1.**
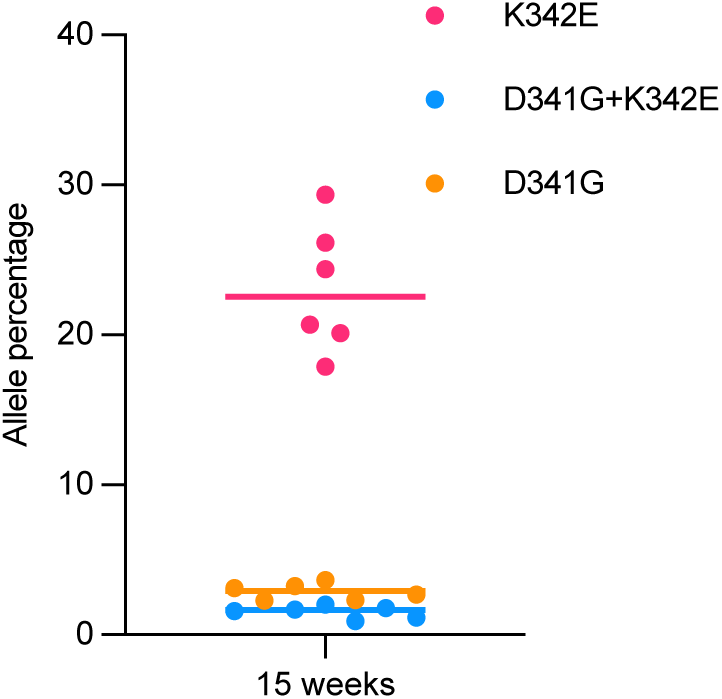
eNme2.C-TadA9e treated AAT-null;PiZ animals allele percentage. AAT-null;PiZ mice were treated with eNme2.C-TadA9e for 15 weeks. Line represents median, n=6.

**Supplemetary Figure S2.**
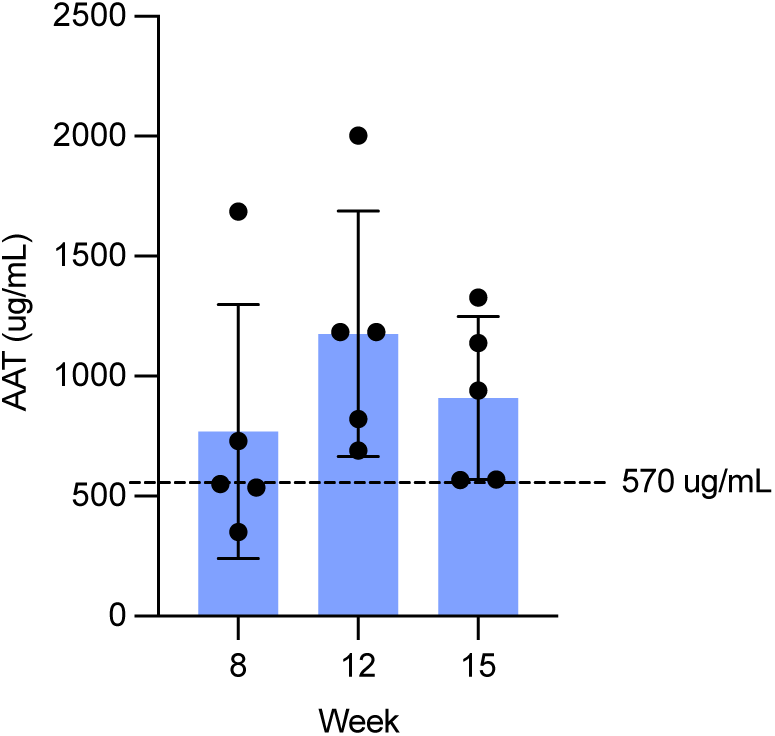
AAT levels in AAT-null;PiZ animals. Z-AAT levels were deducted from total AAT levels in eNme2.C-ABE9e treated animals. Mean ± SD shown, n=5.

## Notes

### Competing Interest Statement

The University of Massachusetts Chan Medical School has filed a patent application on this work.

