## Supplemental file 1 for "A Compact Base Editor Rescues AATD-associated Liver and Lung Disease in Mouse Models"

Supplemental file 1. Off-target sites nominated by Cas-OFFinder

| Guide | | Location | Site | Strand | Mismatches | Notes |
|---|---|---|---|---|---|---|
| CCATCGACAAGAAAGGGACTGAANNNNNNN | chr1 | 1737894 | CCAcCaACAAGgtAGGGACTGgAGGATTCC | - | 5 | |
| CCATCGACAAGAAAGGGACTGAANNNNNNN | chr1 | 15582655 | aCActGAtAAGAAAGGGACTtAATTCTTCC | + | 5 | |
| CCATCGACAAGAAAGGGACTGAANNNNNNN | chr1 | 91298301 | CagTCGAgAAGcAAGGGACTGAgGAGCTCA | + | 5 | |
| CCATCGACAAGAAAGGGACTGAANNNNNNN | chr1 | 159278692 | CCATCcAttAGAAAGaGACTaAACCATACA | + | 5 | |
| CCATCGACAAGAAAGGGACTGAANNNNNNN | chr1 | 187866475 | CCATtGtCAAGAAAGtcACTGAAATATTCT | + | 4 | |
| CCATCGACAAGAAAGGGACTGAANNNNNNN | chr1 | 213693869 | aCATgGgtAAGAAAGGGACTtAAAAGGGCT | + | 5 | |
| CCATCGACAAGAAAGGGACTGAANNNNNNN | chr1 | 223290302 | CCATCGACcAcccAaGGACTGAAGAGCACA | - | 5 | |
| CCATCGACAAGAAAGGGACTGAANNNNNNN | chr10 | 24082082 | aCATacACAgGAAcGGGACTGAAGCGCGCA | - | 5 | |
| CCATCGACAAGAAAGGGACTGAANNNNNNN | chr10 | 29309546 | aCATatgCAAGAAAGGGAaTGAAACATGCT | - | 5 | |
| CCATCGACAAGAAAGGGACTGAANNNNNNN | chr10 | 86448930 | CCATaGACAgaAAtGGGACaGAACAGCACC | - | 5 | |
| CCATCGACAAGAAAGGGACTGAANNNNNNN | chr10 | 105830210 | CCATCccCAAGAAAGGcACTGtgGAGACCA | - | 5 | |
| CCATCGACAAGAAAGGGACTGAANNNNNNN | chr10 | 128846156 | CCATCaACAAGAAAatGgCTaAACAAATCA | - | 5 | |
| CCATCGACAAGAAAGGGACTGAANNNNNNN | chr11 | 1391058 | CCtgtGgCAAGAAAGGGACTcAAGACAGCC | + | 5 | |
| CCATCGACAAGAAAGGGACTGAANNNNNNN | chr11 | 5247758 | CCATaGAaAAGAAgGGGAaaGAAAACATCA | + | 5 | |
| CCATCGACAAGAAAGGGACTGAANNNNNNN | chr11 | 15787200 | tCATCtcCAgGAAAGaGACTGAAAGGGACT | + | 5 | |
| CCATCGACAAGAAAGGGACTGAANNNNNNN | chr11 | 35812057 | gCAgaGACAAGAgAGGaACTGAAGACTGCA | - | 5 | |
| CCATCGACAAGAAAGGGACTGAANNNNNNN | chr11 | 37890968 | CtAagGAaAAGAAAaGGACTGAATAGCTCC | + | 5 | |
| CCATCGACAAGAAAGGGACTGAANNNNNNN | chr11 | 70796156 | CCATCtcCAAaAAAaGGgCTGAAGGCAACT | - | 5 | |
| CCATCGACAAGAAAGGGACTGAANNNNNNN | chr11 | 89886333 | CCATCtACAAGtAAaGGAaTaAACTGTGCT | - | 5 | |
| CCATCGACAAGAAAGGGACTGAANNNNNNN | chr11 | 116720837 | agATtGgCAAcAAAGGGACTGAAGTGACCA | - | 5 | OT 1 |
| CCATCGACAAGAAAGGGACTGAANNNNNNN | chr11 | 126001642 | CCAaCaACcAtcAAGGGACTGAATTCTGCC | + | 5 | |
| CCATCGACAAGAAAGGGACTGAANNNNNNN | chr12 | 4398952 | CCAcgGgCAgGAcAGGGACTGAAGTAACCA | + | 5 | OT 2 |
| CCATCGACAAGAAAGGGACTGAANNNNNNN | chr12 | 7256616 | CCAagGACAAaAAAGGGAgaGAAGATGACA | + | 5 | |
| CCATCGACAAGAAAGGGACTGAANNNNNNN | chr12 | 60036283 | CCATCtAtAAGAcAGGtACTGcATTAAGCT | - | 5 | |
| CCATCGACAAGAAAGGGACTGAANNNNNNN | chr12 | 64148948 | tCATCcACAtGAAAaGGAgTGAAGTATGCC | + | 5 | |
| CCATCGACAAGAAAGGGACTGAANNNNNNN | chr12 | 73540228 | CCATCcACAAGccAGtGtCTGAAGCTACCA | - | 5 | |
| CCATCGACAAGAAAGGGACTGAANNNNNNN | chr12 | 79069032 | CCAggtAtAAGAAAGGcACTGAAAATATCT | + | 5 | |
| CCATCGACAAGAAAGGGACTGAANNNNNNN | chr12 | 94220792 | CCAggGACAAGAAgGGGACaaAATAAACCC | + | 5 | |
| CCATCGACAAGAAAGGGACTGAANNNNNNN | chr12 | 121745699 | CCAaCatCtAGAAAGGGACTGAAAAATGCT | + | 4 | |
| CCATCGACAAGAAAGGGACTGAANNNNNNN | chr13 | 24596812 | CCATCtACAgGAAgtGGAaTGAATTTAGCT | + | 5 | |
| CCATCGACAAGAAAGGGACTGAANNNNNNN | chr13 | 25945806 | aCtgCGAaAAGAAAGGGACTGgATGAGTCC | + | 5 | |
| CCATCGACAAGAAAGGGACTGAANNNNNNN | chr13 | 30380986 | CCATaGACAcaAAAGctACTGAAGAAAGCA | - | 5 | |
| CCATCGACAAGAAAGGGACTGAANNNNNNN | chr13 | 36275993 | aCgaaGACAAGAAAtGGACTGAAAAGCTCC | - | 5 | |
| CCATCGACAAGAAAGGGACTGAANNNNNNN | chr13 | 44171451 | CCATgGcCAAGAAAGGGAaacAAAGGGCCA | + | 5 | |
| CCATCGACAAGAAAGGGACTGAANNNNNNN | chr13 | 88968236 | CtATCGAaAAGAAAGGaAtTcAACCCTACT | - | 5 | |
| CCATCGACAAGAAAGGGACTGAANNNNNNN | chr14 | 33849518 | CaAgCaACAAGAAAGGcACTGtAGGGATCC | + | 5 | |
| CCATCGACAAGAAAGGGACTGAANNNNNNN | chr14 | 43475064 | CaAaCtACAAcAgAGGGACTGAAATAAACG | - | 5 | |
| CCATCGACAAGAAAGGGACTGAANNNNNNN | chr14 | 98429988 | CCAataACAAGcAAGGaACTGAAACCTTCA | - | 5 | |
| CCATCGACAAGAAAGGGACTGAANNNNNNN | chr15 | 74940016 | CCAagGtaAAGAAAGGGACTGAtACATTCT | + | 5 | |
| CCATCGACAAGAAAGGGACTGAANNNNNNN | chr15 | 80447139 | CCATCaACcAGAAAtGGAagGAAGGCGTCT | - | 5 | |
| CCATCGACAAGAAAGGGACTGAANNNNNNN | chr15 | 93135124 | CCATtGACAAGAAAtGcttTGAAATGTACT | + | 5 | |
| CCATCGACAAGAAAGGGACTGAANNNNNNN | chr16 | 68270819 | CCAcCGtCcAGAAAGGGACgaAAGCAGGCA | - | 5 | |
| CCATCGACAAGAAAGGGACTGAANNNNNNN | chr16 | 69239470 | CaATgGACAAGAgAGGGAagGAAAGAAACT | - | 5 | |
| CCATCGACAAGAAAGGGACTGAANNNNNNN | chr16 | 77421688 | CCATCGACAAGActGtGAgTaAAGCAACCT | + | 5 | |
| CCATCGACAAGAAAGGGACTGAANNNNNNN | chr16 | 77471081 | aCATCacCAAGcAAcGGACTGAATAAAACA | + | 5 | |
| CCATCGACAAGAAAGGGACTGAANNNNNNN | chr16 | 80263507 | CCAgCGACttGAAAGtGACTcAATGAATCT | + | 5 | |
| CCATCGACAAGAAAGGGACTGAANNNNNNN | chr17 | 6318249 | CCATCtACAAGccAcGGcCTGAAGCTACCG | + | 5 | |
| CCATCGACAAGAAAGGGACTGAANNNNNNN | chr17 | 11970626 | CCtatGACAAGAAAaGGgCTGAAAATTACC | - | 5 | |
| CCATCGACAAGAAAGGGACTGAANNNNNNN | chr17 | 12810452 | tCtTCaAaAAGAAAGtGACTGAACTACACC | - | 5 | |
| CCATCGACAAGAAAGGGACTGAANNNNNNN | chr17 | 36084649 | CCATaGACAtGgAAGtGAaTGAACATCACT | - | 5 | |
| CCATCGACAAGAAAGGGACTGAANNNNNNN | chr17 | 80407549 | CCtcCacCAAGAAAGGaACTGAAAGTCCCA | + | 5 | |
| CCATCGACAAGAAAGGGACTGAANNNNNNN | chr18 | 22888782 | CCATCcACAAGtAAtGaAaTGAATAGCACC | - | 5 | |
| CCATCGACAAGAAAGGGACTGAANNNNNNN | chr18 | 34192002 | aCtTgaACAAGAAAGGGACTGAcCTGGACT | + | 5 | |
| CCATCGACAAGAAAGGGACTGAANNNNNNN | chr19 | 33315799 | CCgTCaACcAaAAAGGGACTGAgGGAGTCT | - | 5 | |
| CCATCGACAAGAAAGGGACTGAANNNNNNN | chr19 | 48148444 | CCcTCcAgAAtgAAGGGACTGAAAGAGACC | - | 5 | |
| CCATCGACAAGAAAGGGACTGAANNNNNNN | chr19 | 56220924 | aCAatGACAcGAAAGGaACTGAAAATAGCG | - | 5 | |
| CCATCGACAAGAAAGGGACTGAANNNNNNN | chr2 | 3160801 | CCATCGtCAAGAAAttcACTtAAAGCTCCT | + | 5 | |
| CCATCGACAAGAAAGGGACTGAANNNNNNN | chr2 | 10712248 | gCATCGACAAGAAAGGaAaaGAcTCCGTCG | - | 5 | |
| CCATCGACAAGAAAGGGACTGAANNNNNNN | chr2 | 53422423 | CCATgGgCtAGAAgGGaACTGAAGGCCACT | - | 5 | |
| CCATCGACAAGAAAGGGACTGAANNNNNNN | chr2 | 58373257 | CCATtGAtgAGAAAGGGtCTGgATCTTGCC | + | 5 | |
| CCATCGACAAGAAAGGGACTGAANNNNNNN | chr2 | 61035189 | CCATgGACAAGAgAGGGAgaGgAAAAGACA | + | 5 | |
| CCATCGACAAGAAAGGGACTGAANNNNNNN | chr2 | 87772658 | CaAcCtACAAGAAgGaGACTGAACCCACCA | + | 5 | OT 3 |
| CCATCGACAAGAAAGGGACTGAANNNNNNN | chr2 | 112234628 | CaAcCtACAAGAAgGaGACTGAACCCACCA | - | 5 | OT 4 |
| CCATCGACAAGAAAGGGACTGAANNNNNNN | chr2 | 119106445 | ggATgGAaAgGAAAGGGACTGAAATAGACT | + | 5 | |
| CCATCGACAAGAAAGGGACTGAANNNNNNN | chr2 | 122319531 | CaATttACAAGAAAGGGgCTcAAAACTACA | + | 5 | |
| CCATCGACAAGAAAGGGACTGAANNNNNNN | chr2 | 124122684 | aCtTCcAgAAGAAAaGGACTGAAGCTCACG | + | 5 | |
| CCATCGACAAGAAAGGGACTGAANNNNNNN | chr2 | 129687578 | CCgcCGgCAAGcAAGGaACTGAAGCCCACT | - | 5 | |
| CCATCGACAAGAAAGGGACTGAANNNNNNN | chr2 | 139560956 | CaATtGACAtGAAgGGGACTGAgGGACTCA | - | 5 | |
| CCATCGACAAGAAAGGGACTGAANNNNNNN | chr2 | 175418910 | CCAatGAaAAGAAtGtGACTGAAAGAGTCA | + | 5 | |
| CCATCGACAAGAAAGGGACTGAANNNNNNN | chr2 | 200725437 | CCAaCaACAAGgAAGtGAaTGAATAGGTCA | + | 5 | |
| CCATCGACAAGAAAGGGACTGAANNNNNNN | chr2 | 205156127 | aCATCcAaAAGAAAtGGAgTGAAGCAATCA | + | 5 | |
| CCATCGACAAGAAAGGGACTGAANNNNNNN | chr2 | 217499987 | CgAggGAgAAGAAAGGGACTGAgAGGCTCC | - | 5 | |
| CCATCGACAAGAAAGGGACTGAANNNNNNN | chr2 | 232081244 | CCAgCGACAcGActGGGACaGAAGGAGCCT | + | 5 | |
| CCATCGACAAGAAAGGGACTGAANNNNNNN | chr2 | 236714932 | CCATtGAaAAGtAAtGGAgTGAAGTTGTCA | + | 5 | |
| CCATCGACAAGAAAGGGACTGAANNNNNNN | chr2 | 236882794 | CCcTgGACAAGAAAGccACTtAACCTCTCT | - | 5 | |
| CCATCGACAAGAAAGGGACTGAANNNNNNN | chr2 | 241060098 | aCATCGAaAAGAAAGGGAgaaAAACAAACT | - | 5 | |

| | | | | | | |
|---|---|---|---|---|---|---|
| CCATCGACAAGAAAGGGACTGAANNNNNN | chr2 | 241292616 | CCAcaGACAgGAcAGGGACTGAgAATCCCA | + | 5 | |
| CCATCGACAAGAAAGGGACTGAANNNNNN | chr20 | 2960029 | gCATaGAaAAGAAAGGaACTGgATAGGTCC | - | 5 | |
| CCATCGACAAGAAAGGGACTGAANNNNNN | chr20 | 14631226 | CCATaGACAAcAAAtaGgCTGAAAAAGTCA | - | 5 | |
| CCATCGACAAGAAAGGGACTGAANNNNNN | chr21 | 36527693 | CCtTCGtaAAcAAAGGGAgTGAAGTCCCCT | - | 5 | |
| CCATCGACAAGAAAGGGACTGAANNNNNN | chr21 | 47141663 | CCATaGACAgGgAgGGGACTGAgTCTTACT | + | 5 | |
| CCATCGACAAGAAAGGGACTGAANNNNNN | chr22 | 48250891 | CCAcCtACAAGccAGGaACTGAACCCTACG | + | 5 | |
| CCATCGACAAGAAAGGGACTGAANNNNNN | chr3 | 15019102 | CCATaGgaAAGAAAGGaACTaAAAGAAACA | - | 5 | |
| CCATCGACAAGAAAGGGACTGAANNNNNN | chr3 | 15631101 | CCAatcACAAGcAAGGGcCTGAAACAATCA | + | 5 | |
| CCATCGACAAGAAAGGGACTGAANNNNNN | chr3 | 72962890 | gCATgGACAAGAAAGtGACgGtAGTATACA | - | 5 | |
| CCATCGACAAGAAAGGGACTGAANNNNNN | chr3 | 80116990 | CCAattACtAGAAAGGGAaTGAATAAGTCA | - | 5 | |
| CCATCGACAAGAAAGGGACTGAANNNNNN | chr3 | 113125888 | CtAcaaACAAGAAAGaGACTGAATGTACCT | + | 5 | OT 5 |
| CCATCGACAAGAAAGGGACTGAANNNNNN | chr3 | 126169863 | CCATgGAaAgGAAAGGGACTcAAGGCCACT | + | 4 | |
| CCATCGACAAGAAAGGGACTGAANNNNNN | chr3 | 162935666 | gCATCtACAAcAAActGACTGAAAATGGCT | - | 5 | |
| CCATCGACAAGAAAGGGACTGAANNNNNN | chr3 | 194529980 | CCtTtGAtAAGAAAGGGgCTaAATTGCTCA | + | 5 | |
| CCATCGACAAGAAAGGGACTGAANNNNNN | chr4 | 4002927 | CCAcCaACtAaAAAaGGACTGAACAATACT | - | 5 | |
| CCATCGACAAGAAAGGGACTGAANNNNNN | chr4 | 17475539 | CCAgaGcCAtGtAAGGGACTGAAACTGGCG | - | 5 | |
| CCATCGACAAGAAAGGGACTGAANNNNNN | chr4 | 36501884 | CaATaGACAAGAAAtGGAaTcAAGTATCCA | - | 5 | |
| CCATCGACAAGAAAGGGACTGAANNNNNN | chr4 | 46020316 | CtATaGACAAaAAAGGGACTtgAGTGTTCT | + | 5 | |
| CCATCGACAAGAAAGGGACTGAANNNNNN | chr4 | 86858755 | gCATaGtCAAGAAAGaGACTGAcAAGGGCA | + | 5 | |
| CCATCGACAAGAAAGGGACTGAANNNNNN | chr4 | 97233884 | CCATCtAtAAGAAAtGtAaTGAAGGTTACC | - | 5 | |
| CCATCGACAAGAAAGGGACTGAANNNNNN | chr4 | 101570814 | CCATgGgCAAGAAAGGagtTGAATTTAACC | - | 5 | |
| CCATCGACAAGAAAGGGACTGAANNNNNN | chr4 | 137565507 | CCAggtACAAGAAAGGGACTccAACCTCCC | + | 5 | |
| CCATCGACAAGAAAGGGACTGAANNNNNN | chr4 | 138637144 | CCATCatCAAGAAAGtGACTtAgCTGCTCT | - | 5 | |
| CCATCGACAAGAAAGGGACTGAANNNNNN | chr4 | 168544271 | CCATtGACAgGtAAGGGAaTGAcAGGGTCC | - | 5 | |
| CCATCGACAAGAAAGGGACTGAANNNNNN | chr4 | 171588401 | CaAagGACAAGAAgGGGACaGAAAAGCTCT | - | 5 | |
| CCATCGACAAGAAAGGGACTGAANNNNNN | chr5 | 725099 | CCATCtACAgGAAgGGtcCTGAAGGGTGCT | - | 5 | |
| CCATCGACAAGAAAGGGACTGAANNNNNN | chr5 | 809413 | CCATCtACAgGAAgGGtcCTGAAGGGTGCT | - | 5 | |
| CCATCGACAAGAAAGGGACTGAANNNNNN | chr5 | 111450097 | CaATCGAgAAaAAAGGGgCTGgAGTATTCA | - | 5 | |
| CCATCGACAAGAAAGGGACTGAANNNNNN | chr5 | 147934713 | CCATaGACAAaAAAGcGACctAAAACAGCC | - | 5 | |
| CCATCGACAAGAAAGGGACTGAANNNNNN | chr5 | 148674753 | CaAagGAgAAGAAAGGGACTGAATGCATCT | - | 4 | |
| CCATCGACAAGAAAGGGACTGAANNNNNN | chr5 | 161124287 | CaATaGAgcAGgAAGGGACTGAAGCAAACC | + | 5 | |
| CCATCGACAAGAAAGGGACTGAANNNNNN | chr5 | 164951973 | CCATCcACAcGAActGGAgTGAAAGTACCT | - | 5 | |
| CCATCGACAAGAAAGGGACTGAANNNNNN | chr6 | 31081949 | aCATtGACcAGAAAGGGAtTGAATCACCCT | - | 4 | |
| CCATCGACAAGAAAGGGACTGAANNNNNN | chr6 | 32077777 | gCAgtGACAgGAgAGGGACTGAACAGAACA | - | 5 | |
| CCATCGACAAGAAAGGGACTGAANNNNNN | chr6 | 58547296 | tCtTCctCAgGAAAGGGACTGAAAGTCACT | + | 5 | |
| CCATCGACAAGAAAGGGACTGAANNNNNN | chr6 | 89886293 | CCATCtACAAGccAGGGAgaGAAGCTTCCA | + | 5 | |
| CCATCGACAAGAAAGGGACTGAANNNNNN | chr6 | 116269385 | CCAaCaACcAGAAAGGaACTGAtTTCCTCA | + | 5 | |
| CCATCGACAAGAAAGGGACTGAANNNNNN | chr6 | 121758975 | gtAaCGACAAGAAAGatACTGAATATCGCC | + | 5 | |
| CCATCGACAAGAAAGGGACTGAANNNNNN | chr6 | 133364486 | tCATatACAAGcAAGGGAtTGAATTTTACA | + | 5 | |
| CCATCGACAAGAAAGGGACTGAANNNNNN | chr6 | 137285777 | CCAatcACAAGcAAGGGACTGAAATACCCC | - | 4 | |
| CCATCGACAAGAAAGGGACTGAANNNNNN | chr6 | 159197429 | CCATCGACAAGAAgGcacCTGtAAGTTCCA | - | 5 | |
| CCATCGACAAGAAAGGGACTGAANNNNNN | chr6_cox_hap2 | 2596653 | aCATtGACcAGAAAGGGAtTGAATCACCCT | - | 4 | |
| CCATCGACAAGAAAGGGACTGAANNNNNN | chr6_cox_hap2 | 3548439 | gCAgtGACAgGAgAGGGACTGAACAGAACA | - | 5 | |
| CCATCGACAAGAAAGGGACTGAANNNNNN | chr6_dbb_hap3 | 2379121 | aCATtGACcAGAAAGGGAtTGAATCACCCT | - | 4 | |
| CCATCGACAAGAAAGGGACTGAANNNNNN | chr6_dbb_hap3 | 3356951 | gCAgtGACAgGAgAGGGACTGAACAGAACA | - | 5 | |
| CCATCGACAAGAAAGGGACTGAANNNNNN | chr6_mann_hap4 | 2430306 | aCATtGACcAGAAAGGGAtTGAATCACCCT | - | 4 | |
| CCATCGACAAGAAAGGGACTGAANNNNNN | chr6_mcf_hap5 | 2463874 | aCATtGACcAGAAAGGGAtTGAATCACCCT | - | 4 | |
| CCATCGACAAGAAAGGGACTGAANNNNNN | chr6_mcf_hap5 | 3457605 | gCAgtGACAgGAgAGGGACTGAACAGAACA | - | 5 | |
| CCATCGACAAGAAAGGGACTGAANNNNNN | chr6_qbl_hap6 | 2377767 | aCATtGACcAGAAAGGGAtTGAATCACCCT | - | 4 | |
| CCATCGACAAGAAAGGGACTGAANNNNNN | chr7 | 10591500 | tCATaGACAAGAAAGacAtTGAAGCTACCT | + | 5 | |
| CCATCGACAAGAAAGGGACTGAANNNNNN | chr7 | 41328583 | tCATCcACAAGAAAGGGgCTatAGGTCCCA | - | 5 | |
| CCATCGACAAGAAAGGGACTGAANNNNNN | chr7 | 43192924 | aCATCGACAgGAAAGGGAaTacATATGCCC | - | 5 | |
| CCATCGACAAGAAAGGGACTGAANNNNNN | chr7 | 45145660 | CCATaGACAgGgcAGGcACTGAATTTGACA | + | 5 | |
| CCATCGACAAGAAAGGGACTGAANNNNNN | chr7 | 67246667 | taATaGACAAGAAAGGGAgTaAACCTGCCT | + | 5 | |
| CCATCGACAAGAAAGGGACTGAANNNNNN | chr7 | 68885003 | gaAaCGAaAAaAAAGGGACTGAAAAACTCA | + | 5 | |
| CCATCGACAAGAAAGGGACTGAANNNNNN | chr7 | 91054738 | CtATaGtCAAGAAAGGGgCaGAACTTGGCC | - | 5 | |
| CCATCGACAAGAAAGGGACTGAANNNNNN | chr7 | 99130401 | CCtTtGAaAAGAAAtGGACTtAAAGTATCT | + | 5 | |
| CCATCGACAAGAAAGGGACTGAANNNNNN | chr7 | 128410767 | CCATCtgtAAGAAAGGGACccAAAAGGACA | - | 5 | |
| CCATCGACAAGAAAGGGACTGAANNNNNN | chr8 | 26517058 | CtgTCcACAAGAAAGGGACaaAAAGTCTCA | + | 5 | |
| CCATCGACAAGAAAGGGACTGAANNNNNN | chr8 | 33112894 | CaATtGACAAGAAgGGGAgTGtATTGGTCT | + | 5 | |
| CCATCGACAAGAAAGGGACTGAANNNNNN | chr8 | 60357086 | atATtGACAAtAAAtGGACTGAACTCTCCA | - | 5 | |
| CCATCGACAAGAAAGGGACTGAANNNNNN | chr9 | 107816303 | CCATtGACAAGcAAGGcACTGgAGACAGCC | - | 4 | |
| CCATCGACAAGAAAGGGACTGAANNNNNN | chr9 | 109321034 | CCATCcACAtGgAAaGGtCTGAACCAGTCC | - | 5 | |
| CCATCGACAAGAAAGGGACTGAANNNNNN | chr9 | 117875772 | aCtTCGACAAGAAtGtGtCTGAAATGTTCT | - | 5 | |
| CCATCGACAAGAAAGGGACTGAANNNNNN | chr9 | 119137391 | tCATCcACAAGAtAGGtACTGAAAACATCA | - | 4 | |
| CCATCGACAAGAAAGGGACTGAANNNNNN | chrX | 1041124 | CCATCcACAtGgAAGGGACTGgATGGACCA | + | 4 | |
| CCATCGACAAGAAAGGGACTGAANNNNNN | chrX | 1103504 | CCATCcACAtGgAAGGGACTGgATGGACCA | + | 4 | |
| CCATCGACAAGAAAGGGACTGAANNNNNN | chrX | 7126063 | gaATCaACAAcAAAGaGACTGAAATGCTCA | + | 5 | |
| CCATCGACAAGAAAGGGACTGAANNNNNN | chrX | 7084400 | CCAcCGACAAaAAtGGGACTcAAAAATACC | - | 4 | |
| CCATCGACAAGAAAGGGACTGAANNNNNN | chrX | 96128077 | CCATCcAaAAGAAAGGcAaTGAcAAATGCT | - | 5 | |
| CCATCGACAAGAAAGGGACTGAANNNNNN | chrX | 103298864 | gtAcCGgCAAGAAAGGGACTGAcTGAGGCA | - | 5 | |
| CCATCGACAAGAAAGGGACTGAANNNNNN | chrX | 125769563 | CCATtGAatAtAAAtGGACTGAACTCTCCA | - | 5 | |
| CCATCGACAAGAAAGGGACTGAANNNNNN | chrY | 991124 | CCATCcACAtGgAAGGGACTGgATGGACCA | + | 4 | |
| CCATCGACAAGAAAGGGACTGAANNNNNN | chrY | 1053504 | CCATCcACAtGgAAGGGACTGgATGGACCA | + | 4 | |
