## Supplemental file 2 for "A Compact Base Editor Rescues AATD-associated Liver and Lung Disease in Mouse Models"

| Supplemental file 2. Serum AAT levels | | | | |
|---|---|---|---|---|
| Animal number | Treatment | Total AAT ug/ml (ELISA) | | |
| | | 8 weeks | 12 weeks | 15 weeks |
| 912 | Control | 392.9 | 241.4 | 212.4 |
| 945 | | 622.5 | 754.6 | 529 |
| 947 | | 638.1 | 924.5 | 736.2 |
| 922 | ABE | 902.1 | 1329.9 | 1063.2 |
| 917 | | 1086.9 | 1461.7 | 1059.7 |
| 950 | | 1101.4 | 1824.5 | 1433.8 |
| 951 | | 1280.6 | 1824.3 | 1632 |
| 921 | | 2237.5 | 2643.2 | 1821.3 |
