## Supplemental file 3 for "A Compact Base Editor Rescues AATD-associated Liver and Lung Disease in Mouse Models"

| Supplemental File 3. Primers used for high-throughput sequencing | | |
|---|---|---|
| Site | Forward primer | Reverse primer |
| PiZ reporter cell | CTACACGACGCTCTTCCGATCTAAGGTCTTCAGCAATGGGGC | AGACGTGTGCTCTTCCGATCTAGGGTTTGTTGAACTTGACCTC |
| *SERPINA1* | CTACACGACGCTCTTCCGATCTAACGTGTCTCTGCTTCTCTCC | AGACGTGTGCTCTTCCGATCTGGGATTCACCACTTTTCCCATGAAGAGGGGAGAC |
| OT1 | CTACACGACGCTCTTCCGATCTAGGGGATTGGTTATGAGGCT | AGACGTGTGCTCTTCCGATCTGCTCTTGGGGCCACTATTGAT |
| OT2 | CTACACGACGCTCTTCCGATCTAAGAGGTCAGAGAGGCCATT | AGACGTGTGCTCTTCCGATCTAAGGGTGGAGTGGGAAGTTT |
| OT3 | CTACACGACGCTCTTCCGATCTTTGGGGCCAAAAACAAAACA | AGACGTGTGCTCTTCCGATCTCTTGAAAAGAGGCTCTCAGC |
| OT4 | CTACACGACGCTCTTCCGATCTCTTGAAAAGAGGCTCTCAGC | AGACGTGTGCTCTTCCGATCTTTGGGGCCAAAAACAAAACA |
| OT5 | CTACACGACGCTCTTCCGATCTAGACCGGTGAACGTAAAAGC | AGACGTGTGCTCTTCCGATCTTTGCCGGGCAGGTAGTAGAT |
