## Supplemental file 4 for "A Compact Base Editor Rescues AATD-associated Liver and Lung Disease in Mouse Models"

| Supplemental file 4. Guides used for PiZ editing. | | | |
|---|---|---|---|
| Guide | Spacer | PAM | Effector |
| Guide 1 | tgaccatcgacaagaaagggact | gaagct | eNme2.C |
| Guide 2 | ccatcgacaagaaagggactgaa | gctgct | eNme2.C |
